# Single-cell transcriptomic profiling of the neonatal oviduct and uterus reveals new insights into upper Müllerian duct regionalization

**DOI:** 10.1101/2023.12.20.572607

**Authors:** Shuai Jia, Fei Zhao

## Abstract

The upper Müllerian duct (MD) is patterned and specified into two morphologically and functionally distinct organs, the oviduct and uterus. It is known that this regionalization process is instructed by inductive signals from the adjacent mesenchyme. However, the interaction landscape between epithelium and mesenchyme during upper MD development remains largely unknown. Here, we performed single-cell transcriptomic profiling of mouse neonatal oviducts and uteri at the initiation of MD epithelial differentiation (postnatal day 3). We identified major cell types including epithelium, mesenchyme, pericytes, mesothelium, endothelium, and immune cells in both organs with established markers. Moreover, we uncovered region-specific epithelial and mesenchymal subpopulations and then deduced region-specific ligand-receptor pairs mediating mesenchymal-epithelial interactions along the craniocaudal axis. Unexpectedly, we discovered a mesenchymal subpopulation marked by neurofilaments with specific localizations at the mesometrial pole of both the neonatal oviduct and uterus. Lastly, we analyzed and revealed organ-specific signature genes of pericytes and mesothelial cells. Taken together, our study enriches our knowledge of upper Müllerian duct development, and provides a manageable list of potential genes, pathways, and region-specific cell subtypes for future functional studies.

## Introduction

A fundamental question in developmental biology is how a primitive epithelial tube differentiates into morphologically and functionally distinct tubular organs along the craniocaudal axis. During development, the Müllerian duct (MD), the primordium of female reproductive tract organs, undergoes craniocaudal patterning from a simple tube into four distinct organs: the oviduct, uterus, cervix, and vagina (1). During postnatal differentiation, epithelial cells in these organs acquire unique phenotypes and functions. In the upper MD, epithelial cells are committed to become oviductal ciliated and secretory cells (2); in the middle MD, they differentiate into uterine luminal and glandular epithelial cell types (3); in the lower MD, they become basal and stratified squamous epithelium in the ectocervix and vagina (4). Impairment of MD differentiation by genetic and environmental factors can potentially lead to Müllerian anomalies, a major cause of female infertility in humans (5, 6).

MD regionalization is largely dependent on instructive signals from its surrounding mesenchyme. MD mesenchyme expresses region-specific Homeobox genes for controlling MD craniocaudal patterning: *Hoxa9* in the oviduct, *Hoxa10* in the uterus, *Hoxa11* in the posterior uterus and cervix; and *Hoxa13* in the cervix and upper vagina (7, 8). Genetic ablations of these *Hox* genes expressed in the mesenchyme lead to homeotic transformation in mice, providing critical evidence that the mesenchyme governs MD epithelial fate and differentiation. For example, *Hoxa10* knockout caused partial morphological alteration of the uterus to the oviductal appearance(9); when the uterine *Hox* gene *Hoxa11* was swapped with *Hoxa13* that is normally expressed in the mesenchyme of the cervix and vagina, the uterus underwent homeotic transformation and became similar to the cervix and vagina (10). Classic tissue recombination studies also demonstrate the instructive function of the mesenchyme in MD regionalization. Heterotypic tissue recombinants of uterine mesenchyme and vaginal epithelium developed into the uterus while recombinants of vaginal mesenchyme and uterine epithelium underwent vaginal morphogenesis (11, 12). In tissue recombinants, developmental plasticity of epithelial cells is gradually lost approximately by 10 days postpartum, after which epithelial cells cannot be reprogrammed (11, 12). Therefore, region-specific mesenchymal signals at early postnatal development govern MD craniocaudal patterning.

The paracrine signals from the mesenchyme for determining the fate and differentiation of the middle and lower MD have been discovered. WNT4 and WNT5A in the mesenchyme regulates the differentiation of uterine glandular epithelium (Kelleher.et al. 2019). Activin A, BMP4 and FGF7/10 from the lower MD mesenchyme are independently required for inducing vaginal cell fate (Terakawa. et al. 2020). However, it is unknown what mesenchymal factor(s) determines the fate of upper MD epithelium to become oviductal epithelia. In an effort to address this knowledge gap, we performed single-cell mRNA profiling of the neonatal oviduct and uterus to reveal region-specific interaction landscape between epithelium and mesenchyme at postnatal day 3 (PND3), a critical developmental timepoint for establishing regional patterns (11–13).

## Results

### Overview and characterization of major cell populations in the neonatal oviduct and uterus

Our scRNA-seq profiling of PND3 oviducts and uteri generated a dataset with 22150 individual cells after filtering with average 3006 genes/cell (**Supplementary file 1**). Based on their transcriptomic similarities, those individual cells were classified into 8 cell populations with distinct cell-type specific markers: oviductal mesenchymal cells (*Hoxc8,* 7438 cells without expression of proliferation markers, 38%), uterine mesenchymal cells (*Hoxa10* (14), 4898 cells without expression of proliferation markers, 25%), proliferating mesenchymal cells (*Top2a* (15), 2840, 14%), epithelial cells (*Epcam1*(*16*), 1150, 6%), endothelial cells (*Pecam1*(*17*), 1058, 5%), pericytes (*Rgs5*(*18*), 778, 4%), mesothelial cells (*Msln*(*19*)*, 1139, 6%*), and myeloid/immune cells (*Lzy2*(*20*), 424, 2%) (**Fig. 1A-1C**). Through differential expression analysis, we identified the top 5 significantly enriched genes in each cluster **(Fig. 1D**), some of which could serve as new cell type markers at this developmental stage. For example, *Enpp2* for oviductal mesenchymal cells, *Tcf21* for uterine mesenchymal cells (validated in the following figures), and *Wfdc2* for epithelial cells **(Fig. 1D & Supplementary file 2**).

**Figure 1.**
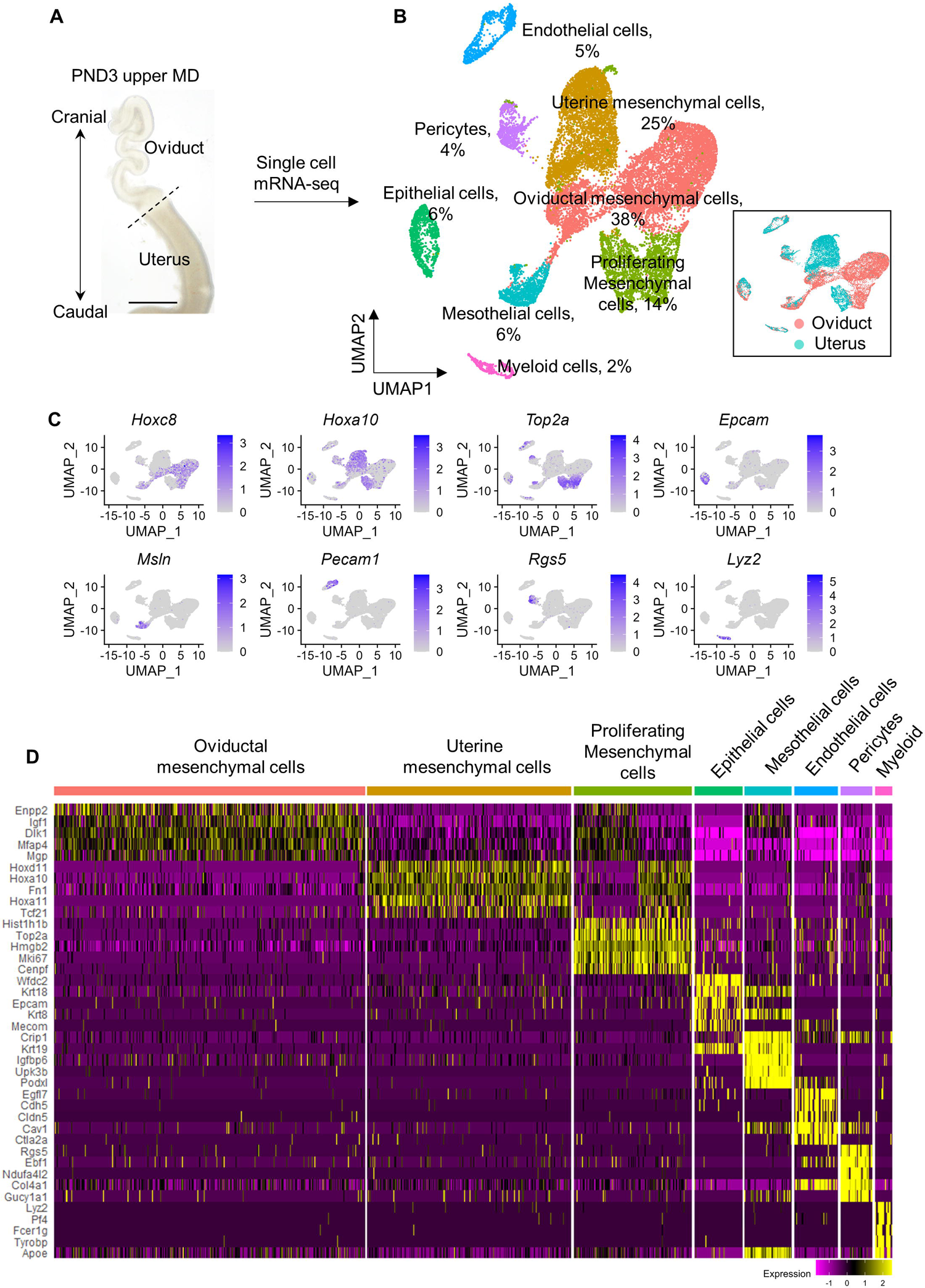
Single-cell transcriptional atlas of PND3 oviduct and uterus. (A) Tissue sample surveyed in this study. The dotted line indicated where the oviduct and the uterus were separated. Scale bar=2.5 mm. (B) Uniform manifold approximation and projection (UMAP) plot featuring general cell clusters from both the oviduct and uterus. In the lower right, the same UMAP was annotated according to cell’s organ origin. (C) UMAPs featuring cell type markers, *Hoxc8* for oviductal mesenchyme, *Hoxa10* for uterine mesenchyme, *Top2a* for proliferating cells, *Epcam* for epithelium, *Msln* for mesothelium, *Pecam1* for endothelium, *Rgs5* for pericytes, and *Lyz2* for myeloid cells. (C) Heatmap of top five marker genes for each cluster.

When UMAP was annotated based on organ origins of each individual cells, both uterine and oviductal cells were overlapped in the clusters of the epithelia, mesothelia, endothelia, pericytes and myeloid cells but separated in two distinct mesenchymal clusters (**Fig. 1B**). These results demonstrate that both neonatal oviducts and uteri are composed of the same cell types but with the largest transcriptomic difference observed in their mesenchymal cells. This observation is consistent with the notion that region-specific mesenchymal signaling plays inductive roles in MD regionalization (11, 12).

### Dissecting mesenchymal cells of the neonatal oviduct and uterus

Given the importance of the mesenchyme in directing epithelial differentiation and morphogenesis (21), we further dissected heterogeneity of the mesenchymal population, which represented the largest population (63% of cells) in the neonatal oviduct and uterus. We isolated and re-classified mesenchymal population into 7 clusters demarcated by distinct marker gene expression and organ origins (**Fig. S1, 2A & 2B; Supplementary file 3**): 2 smooth muscle subpopulations distinguished by the organ-specific expression of *Tcf21* (in the uterus) and *Nrp2* (in the oviduct); 2 mesenchymal subpopulations (*Pkib*+ and *Pcdh10*+) in the uterus; 2 mesenchymal subpopulations (*Wfdc1*+ and *Angptl7*+) in the oviduct; 1 shared population (*Nefm*+) observed in both the uterus and oviduct.

The uterine and oviductal smooth muscle clusters both expressed *Cnn1*, an established smooth muscle marker (22) (**Fig. 2B**). This observation was consistent with the previous report that smooth muscle differentiation has initiated in the prospective circular myometrial layer at PND3 in both organs (22). However, our scRNA-seq analysis uncovered previously unrecognized gene expression differences between the two smooth muscle populations. *Tcf21* is a bHLH transcription factor and plays critical roles in organogenesis (23). Its expression was enriched specifically in the uterine smooth muscle cells (**Fig. 2B & 2C**). On the other hand, the oviducal smooth muscle cells expressed *Nrp*2 which encodes a transmembrane protein Neuropilin 2 (24). Based on the online scRNA-seq dataset of adult murine oviducts (25), *Tcf21* expression was still not detected but *Nrp2* continued to be expressed in the oviductal smooth muscle cells.

**Figure 2.**
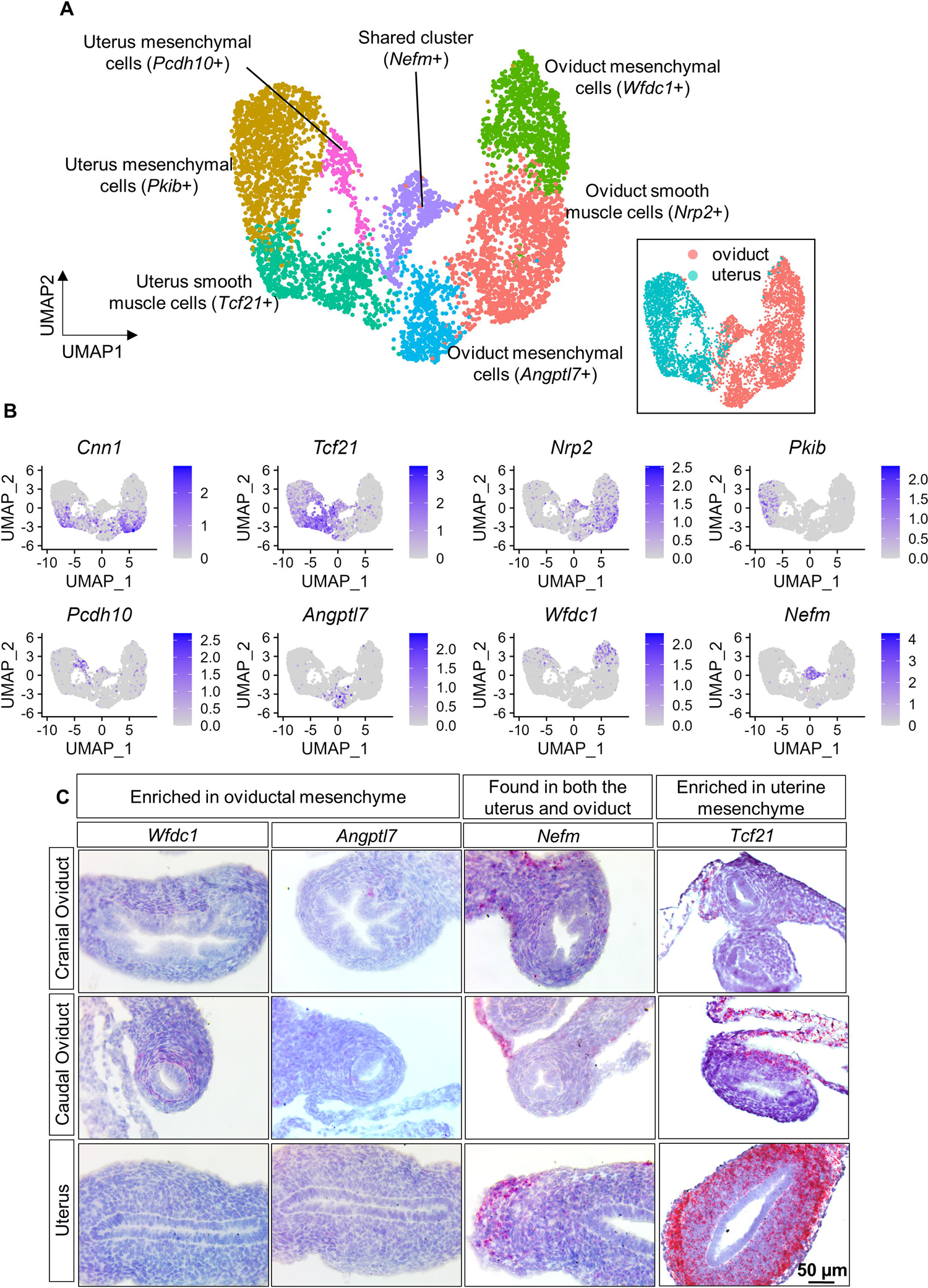
Characterization of mesenchymal subpopulations in PND3 oviduct and uterus. (A) UMAP featuring the mesenchymal subpopulation after re-clustering with higher resolution. In the lower right, the same UMAP was annotated according to cell’s organ origin. (B) UMAPs feature marker genes for cell subtypes: *Cnn1* for smooth muscle cells in both the oviduct and uterus, *Tcf21* for uterine smooth muscle, *Nrp2* for oviductal smooth muscle, *Pkib* & *Pcdh10* for two uterine mesenchymal subtypes, *Wfdc1* & *Angptl7* for two oviductal mesenchymal subtypes, and *Nefm* for a mesenchymal subtype found in both the oviduct and uterus. (C) mRNA expression of four markers for mesenchymal subpopulations, *Wfdc1*, *Angptl7*, *Nefm* and *Tcf21,* by RNAscope. N=3. Scale bar=50 µm.

In addition to smooth muscle cells, there were two mesenchymal subpopulations in either oviducts or uteri. The two mesenchymal subpopulations in oviducts were defined by the expression of *Wfdc1* and *Angptl7*, respectively (**Fig. 2A & 2B**). We validated their expression in the mesenchyme of cranial and caudal oviduct by RNAscope (**Fig. 2C**). At this developmental stage, epithelial cells in the cranial oviduct (future infundibulum and ampulla) undergo longitudinal folding while those in the cauda region (isthmus) remain unfolded (2) (**Fig. 2C**). Both cranial and caudal regions of the oviduct possessed *Wfdc1*+ and *Anpgtl7*+ mesenchymal cells, suggesting that these two populations were not region-specific. However, it seemed that *Wfdc1*+ cells were more enriched in the mesenchyme adjacent to the epithelium of the caudal oviduct, while *Angptl7* were more enriched in the mesenchyme of the cranial oviduct (**Fig. 2C**). Using the online scRNA-seq dataset of adult oviducts (25), we found that *Wfdc1* was still expressed specifically in the mesenchyme-derived fibroblasts in both cranial (infundibulum and ampulla) and cauda (isthmus and uterotubal junction) regions while *Angptl7* expression became undetected. These results demonstrate that *Wfdc1* can serve as a specific marker for oviductal mesenchyme at both neonatal and adult stages while *Angptl7* is a transient marker for oviductal mesenchyme at the neonatal stage.

Contrary to the oviductal mesenchyme, the uterine mesenchyme was marked by the expression of *Tcf21* and classified into *Pkib*+ and *Pcdh10*+ subpopulations (**Fig. 2B & 2C**). Previous studies have identified two closely related mesenchymal subpopulations as inner and outer stroma depending on their proximity to the epithelial lumen in rat uteri at PND6 (26). Markers genes for inner stroma (*Plac8* and *Bmp7)* and outer stroma (*Apoe*) were preferentially expressed in *Pkib*+ and *Pcdh10*+ mesenchymal subpopulations, respectively (**Fig.S2**). This observation suggests that two uterine mesenchymal subpopulations at PND3 in our study may correspond to inner and outer stromal populations at a later developmental stage. In the adult uterine mesenchyme, Kirkwood et al., identified three stromal subpopulations, inner stroma, outer stroma, and the third one localized in the subepithelial space (27). One of the markers (*Col6a4*) for the subepithelial stromal subpopulation in the adult uterus was overrepresented in the *Pkib*+ mesenchyme (potentially inner stroma) in our dataset (**Fig. S2**).

Finally, we identified a new mesenchymal subpopulation demarcated by unique expression of *Nefm* and *Nefl* (**Fig. 2B & Fig. S2**). We found that *Nefm* expression was predominantly localized at the mesometrial pole. In the oviduct, *Nefm* was found in the mesosalpinx ligament and mesenchymal cells at the mesometrial pole. Likewise, in the uterus, it was expressed in the mesometrium as well as mesenchymal cells at the mesometrial pole (**Fig. 2C**). Since *Nefm* (encode neurofilament medium polypeptide) and *Nefl* (encode neurofilament light polypeptide) are neurofilament genes (28), we initially thought that this mesenchymal subpopulation might represent neuronal cells. However, these neurofilament proteins are also expressed in non-neuron cells (29); and *Rbfox3* (encode NeuN (30), a neuronal nuclei marker protein) was not expressed in this population at all (**Fig. S2**). Therefore, we conclude that the *Nefm*+ subpopulation localized at the mesometrial pole is a unique mesenchymal cell subtype expressing neurofilaments as a part of its cytoskeleton network.

### Dissecting epithelial cells of the neonatal oviduct and uterus

The mature epithelium of the oviduct and uterus consists of four epithelial types: secretory and ciliated cells in the oviduct (31, 32); and luminal and glandular epithelial cells in the uterus (3). Establishment of these cell types from MD nascent epithelium does not occur until PND4 (2, 33–35). When comparing epithelial cells from the uterus and oviduct (**Fig. 3A**), we identified 390 differentially expressed genes (DEGs) and the top marker *Msx1* for uterine epithelium and *Lrpap1* for oviductal epithelium (**Fig. 3B and Supplementary file 4**). We confirmed the specific expression of MSX1 in the uterine epithelium but not in the oviductal epithelium on PND3 (**Fig. 3E**). It has been reported that MSX1 is present exclusively in lower parts of the female reproductive tract (the uterus, cervix and vagina) (36). *Msx1* expression continues to be absent in the oviduct at adulthood based on the published scRNA-seq dataset (25). Therefore, we concluded that *Msx1* represents a specific gene signature of postnatal uterine epithelium.

**Figure 3.**
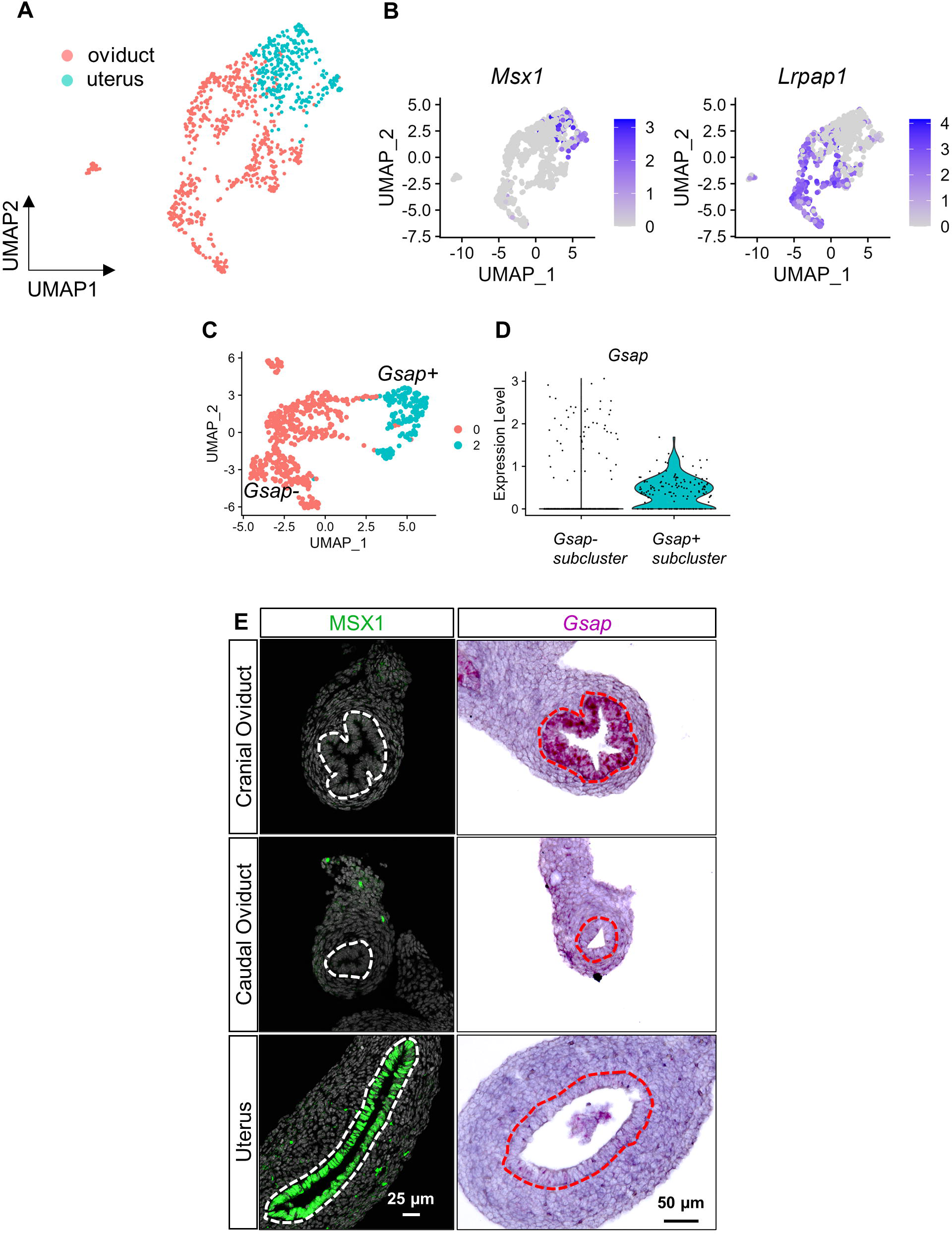
Region-specific epithelial subtypes in PND3 oviduct and uterus. (A) UMAP featuring oviductal and uterine epithelial cells extracted from the overall dataset in Fig 1B. (B) UMAPs featuring marker genes *Msx1* and *Lrpap1* for uterine and oviductal epithelium, respectively. (C) UMAP featuring oviductal epithelial cells extracted from Fig 3A and then re-clustered. (D) Violin plot showing *Gsap* expression in two subclusters in Fig 3C. (E) Immunofluorescent staining of MSX1 and RNAscope staining of *Gsap* in the cranial oviduct, caudal oviduct and uterus at PND3. N=3. Scale bar: 25 and 50 µm for immunofluorescence and RNAscope, respectively.

In the oviduct, morphological differences (epithelial folding vs non-folding) in the cranial and caudal regions were noticed at PND3 (**Fig. 2C**). In addition, it is known that the predominant cell types lining the epithelium of the future cranial and caudal regions are distinct, ciliated epithelial cells enriched in the cranial region while secretory epithelial cells enriched in the caudal region (31). Therefore, we suspect there could be at least two epithelial subtypes in the oviduct at PND3. By increasing the clustering resolution (0.1), we were able to identify *Gsap*+ and *Gsap*-epithelial subtypes at PND3 (**Fig. 3C & 3D**). By performing RNAscope, we discovered that *Gsap* expression was restricted to the cranial oviduct epithelium and barely detected in the caudal oviduct or uterus (**Fig.3E**). Expression of a previously-characterized marker *Wt1* for the cranial oviduct at PND4 (34) was higher in the *Gsap+* population compared to *Gsap-* cluster (**Supplementary file 5**). These results demonstrate the spatial locations of three epithelial subpopulations along the craniocaudal axis at PND3: *Gsap*+ epithelia in the cranial oviduct; *Gsap*-epithelia at the caudal oviduct, and *Msx1*+ uterine epithelia.

In the UMAP clustering, we noticed an outlier distant from the major oviductal epithelial population (**Fig. S3**). When comparing the transcriptome between this outlier and the rest of oviductal cells, we found enrichment of markers for ciliated cells (*Foxj1* and *Fam183b* (37)) and other well-established transcription factors (*Ruvbl1* (*38*) and *Myb* (*39*)) critical for multiciliogenesis, suggesting that this outlier cluster represented ciliated epithelial cells (**Fig. S3**). Therefore, multiciliated cell differentiation has occurred in some of oviductal epithelium at PND3, earlier than previous reports (2, 34). In the list of genes overrepresented in this multiciliated cell cluter (**Fig. S3 and Supplementary file 6**), we uncovered two new transcription factors *Pbx4* and *Aes* (also known as *Tle5*), which might potentially play crucial roles in multiciliated cell differentiation.

### Interrogate ligand-receptor mediated crosstalk between epithelial and mesenchymal cells

After uncovering mesenchymal heterogeneity and epithelial regionalization, we next explored the cell-cell interactions between them by leveraging Cellchat, a bioinformatic tool that quantitatively infers and analyzes ligand-receptor mediated intercellular communication networks (40). The cell-cell interactions between three spatially distinct epithelial subpopulations (*Gsap+* cranial oviductal epithelium, *Gsap-* caudal oviductal epithelium, and *Msx1+* uterine epithelium) and their corresponding mesenchymal populations were analyzed (**Fig. S4 and Supplementary files 7-9**).

First, we inferred the signals sending from the mesenchyme to the epithelium. We identified 157 ligand-receptor pairs in the communication from oviductal mesenchyme to *Gsap*+ cranial oviductal epithelium, 58 from oviductal mesenchyme to *Gsap*-caudal oviductal epithelium and 47 from uterine mesenchyme to uterine epithelium (**Fig. S4 and Supplementary files 10**). In the predicted cell-cell interaction lists, a ligand was paired with multiple receptors/coreceptors (**Supplementary file 7-9**). To concisely visualize the potentially functional ligand-receptor pairs, we focused on mesenchyme-derived ligands for the following criteria: 1) the knockout mice had reproductive phenotypes and/or embryonic lethality based on Mouse Genome Informatics (MGI) (41); 2) Ligands are enriched only in either the oviductal or uterine mesenchyme (**Fig. S4 and Supplementary files 7-9, 11**). Therefore, we depicted a simplified version of mesenchymal-epithelial crosstalk represented by potentially functional region-specific ligands (**Fig. 4**).

**Figure 4.**
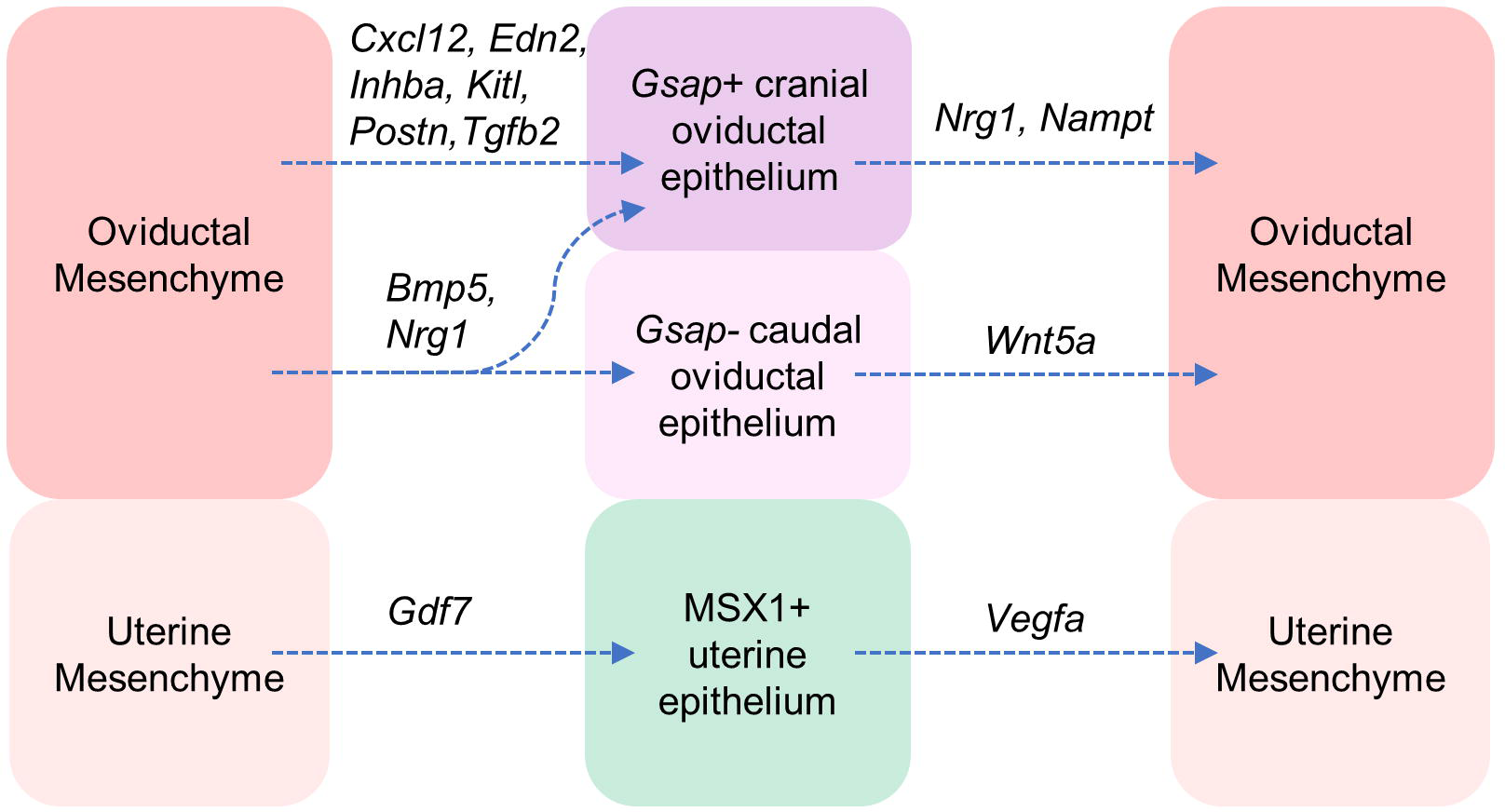
Mesenchymal-epithelial crosstalk represented by region-specific ligands that are potentially functional. Only region-specific ligands whose knockout mice display embryonic lethality and/or reproductive phenotype were included. Please see all ligand-pair pairs in the Fig S4 and supplementary files 7-12.

In the ligand-mediated communication from oviductal mesenchyme to epithelium, *Cxcl12*, *Edn2*, *Inhba*, *Kitl*, *Postn*, *Tgfb2* ligands were specific to *Gsap*+ cranial oviduct while *Bmp5* and *Nrg1* from the mesenchyme can act on both *Gsap*+ and *Gsap*-oviductal epithelium (**Fig. 4**). *Gdf7*, a ligand in the TGF-β pathway, specifically mediated the communication from uterine mesenchyme to epithelium (**Fig. 4**). Although *Gdf7* expression was detected in the uterine stroma (42), its functional significance in uterine differentiation remains to be covered.

The crosstalk between the epithelium and mesenchyme is never a one-way street (12). We therefore inferred the signals sending from the epithelium to the mesenchyme. We identified 144 ligand-receptor pairs from *Gsap*+ oviductal epithelium (cranial oviduct) to oviductal mesenchyme, 70 pairs from *Gsap*-oviductal epithelium (cauda oviduct) to oviductal mesenchyme and 54 ligand-receptor pairs from uterine epithelium to uterine mesenchyme (**Fig. S4, Supplementary file 12**). Likewise, we consolidated and focused on region-specific ligands that were potentially functional (**Supplementary files 7-9,11**). The region-specific ligands deriving from the epithelium to act on adjacent mesenchyme were *Nrg1* and *Nampt* in the cranial oviduct; *Wnt5a* in the caudal oviduct; *Vegfa* in the uterus (**Fig. 4**). *Nrg1* (Neuregulin 1) belongs to the epidermal growth factor and plays essential roles in the heart and nervous system (43). *Nampt* (nicotinamide phosphoribosyltransferase) exists in both intracellular and extracellular forms. The intracellular NAMPT regulates NAD biosynthesis while extracellular NAMPT is proposed to act as an insulin-mimetic cytokine (44, 45). Global mouse knockout of either *Nrg1* or *Nampt* leads to embryonic lethality (43, 44). The availability of *Nrg1-flox* (46) and *Nampt-flox* (47) alleles makes it possible and as a promising direction to investigate their potential roles in MD development. Our finding of *Wnt5a*-mediated communication from the epithelium to the mesenchyme in the caudal oviduct was consistent with the report of enriched *Wnt5a* expression in the epithelium of caudal oviducts at PND28 (33). *Vegfa* have been implicated in embryonic development (48) and adult uterine function (49); however, its specific role in female reproductive tract development remains to be investigated. Taken together, our data provides a list of region-specific ligands/their activated pathways that potentially mediate epithelial-mesenchymal communication during MD regionalization.

### Inquire gene expression differences between oviductal and uterine pericytes and mesothelial cells

Lastly, we turned our attentions to two supporting cells, pericytes (*Rgs5*+ cluster in **Fig. 1**) and mesothelial cells (*Msln*+ cluster in **Fig. 1**), which are rarely studied despite their crucial roles in organ development (50, 51). Pericytes exhibited inter-organ molecular differences (number of DEGs=186, **Supplementary file 13**). The top enriched genes in the oviductal pericytes (*Cebpb*, *Rgs16*, *Nr2f1*, *Hoxc8*, *Cryab*) and uterine pericytes (*Amer1*, *Plac8*, *Hoxd11*, *Lsp1*, *Hoxa10*) were also overrepresented in mesenchymal populations (**Fig. 5A & 5B**), suggesting that pericytes in the oviduct and the uterus might originate from organ-specific mesenchymal cells and/or acquire the organ signature after specification from other cell types (52).

**Figure 5.**
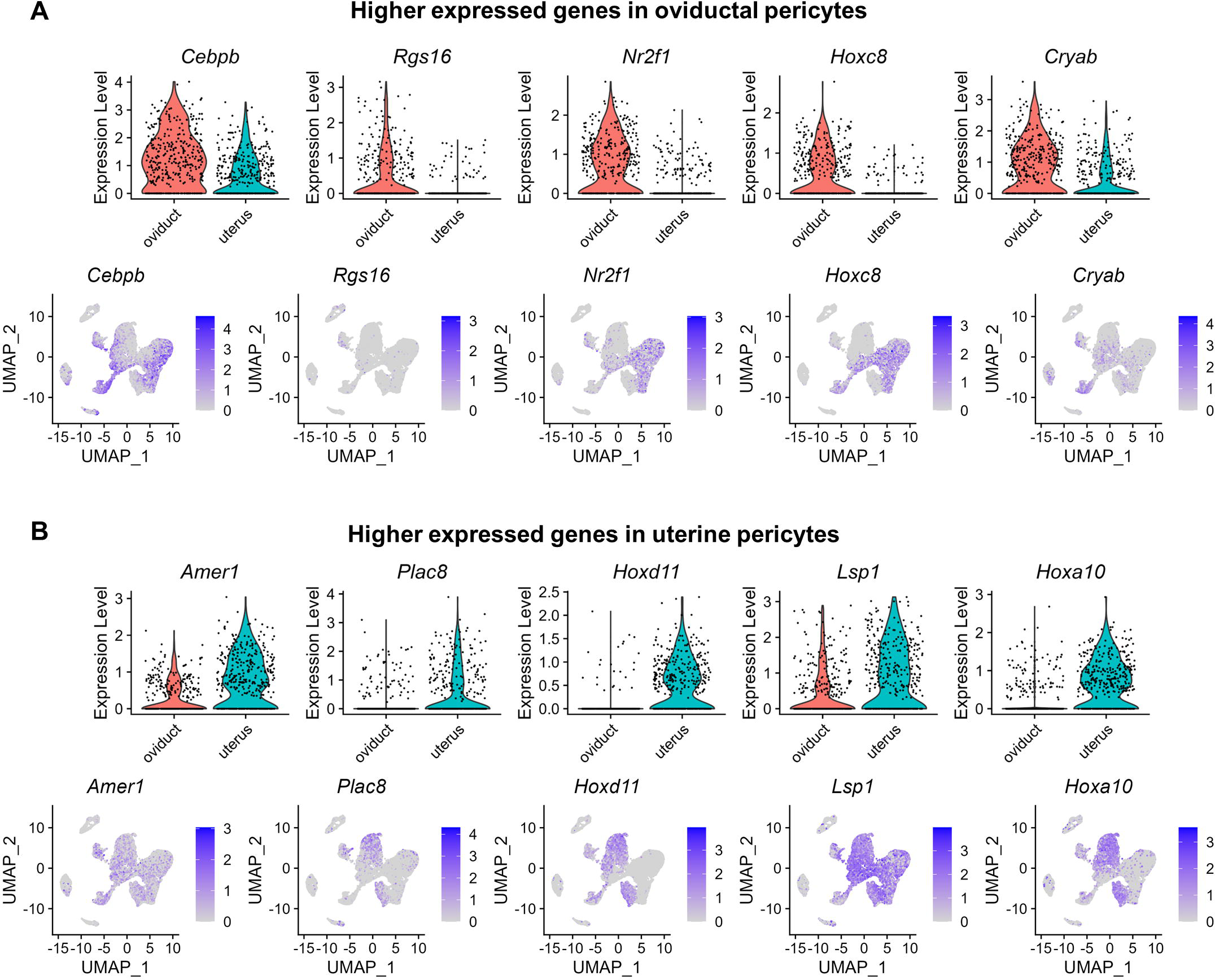
Gene expression comparison between oviductal and uterine pericytes. (A) Violin plots and UMAPs of higher expressed genes (*Cebpb*, *Rgs16*, *Nr2f1*, *Hoxc8* and *Cryab*) in oviductal pericytes. (B) Violin plots and UMAPs of higher expressed genes (*Amer1*, *Plac8*, *Hoxd11*, *Lsp1*, and *Hoxa10*) in uterine pericytes

Mesothelium is an extensive monolayer of squamous-like epithelial cells lining the surface of the oviduct and uterus (53, 54). Both mesothelial and epithelial cells expressed cytokeratins *Krt8*, *Krt18*, and *Krt19* (**Fig. 6A**). However, mesothelial cells did not expression luminal epithelium marker *Epcam* and *Mecom* but instead exhibited the expression of *Msln* (19) and *Upk3b* (55) in both the oviduct and uterus (**Fig. 1B & Fig. 6A**). Transcriptomic analysis between oviductal and uterine mesothelial cells identified 260 DEGs (**Supplementary file 14**), including the top genes enriched in the oviductal mesothelium (*Lgals7*, *Igf1*, *Mt1*, *Ccl2*, and *Nrgn)* and uterine mesothelium (*Alcam*, *Rspo3*, *Fxyd3*, *Tcf21*, and *Hoxd11)* (**Fig. 6B**). It has been reported that *Igf1* is enriched in the cranial MD (future oviduct) (56). Our *Tcf21* RNAscope staining confirmed the enriched expression of *Tcf21* in the mesothelial layer of the uterus but not in the oviduct (**Fig. 2C**). These results validated organ-specific signature genes for mesothelial cells in the neonatal oviduct and uterus.

**Figure 6.**
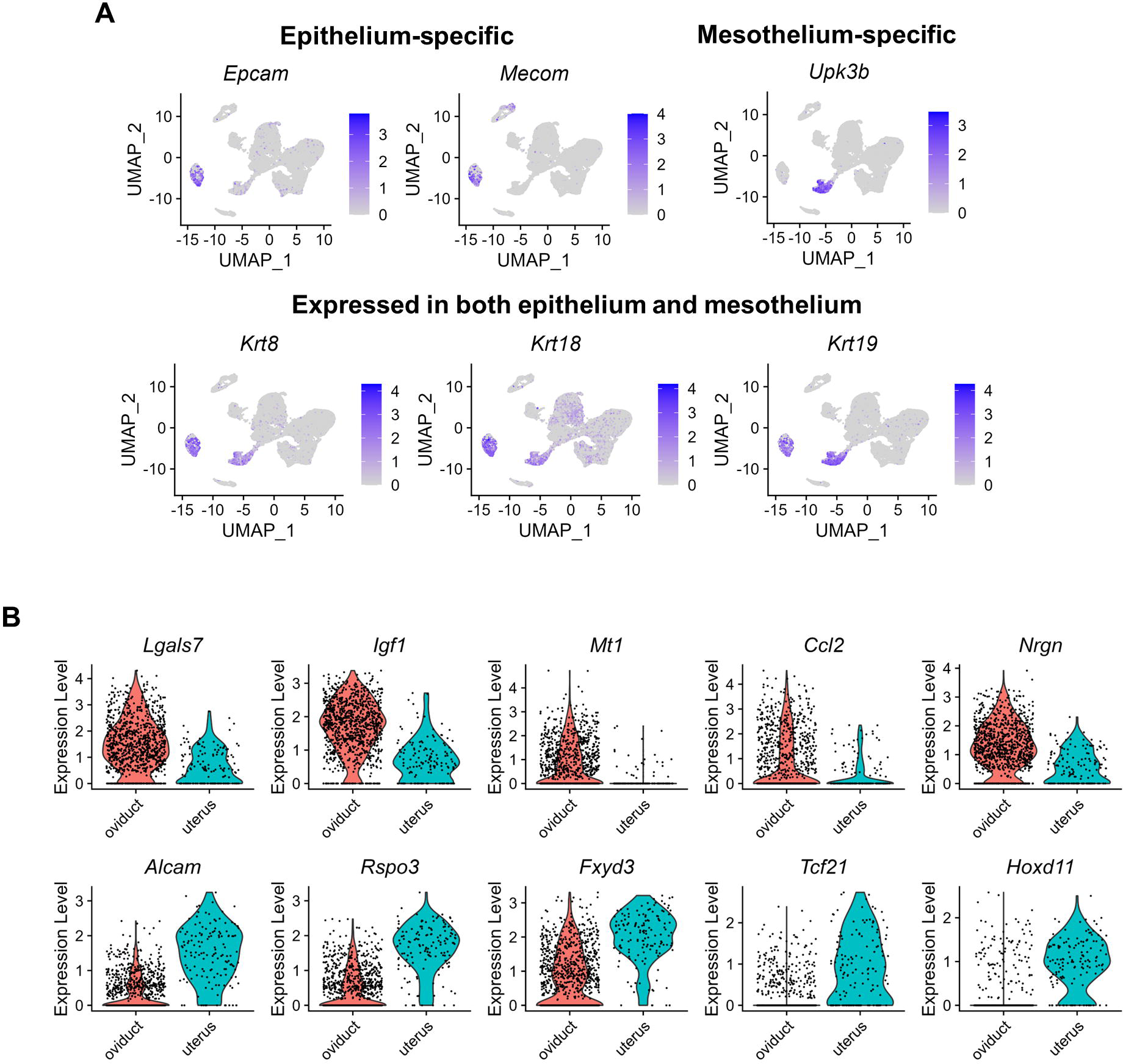
Gene expression comparison between oviductal and uterine mesothelium. (A) UMAPs featuring expression of epithelium-specific (*Epcam*, *Mecom*), mesothelium-specific (*Upk3b*), and commonly shared (*Krt8*, *Krt18* and *Krt19*) marker genes. (B) Violin plots of higher expressed genes in the oviductal mesothelium (up row: *Lgals7*, *Igf1*, *Mt1*, *Ccl2*, and *Nrgn)* and uterine mesothelium (bottom row: *Alcam*, *Rspo3*, *Fxyd3*, *Tcf21*, and *Hoxd11)*.

## Discussion

Our study presented a single-cell atlas of the murine oviduct and uterus at PND3, a critical developmental timepoint for establishing regional differentiation of MD epithelium (11–13). Our major findings on upper MD regionalization were summarized in **Fig. 7**.

**Figure 7.**
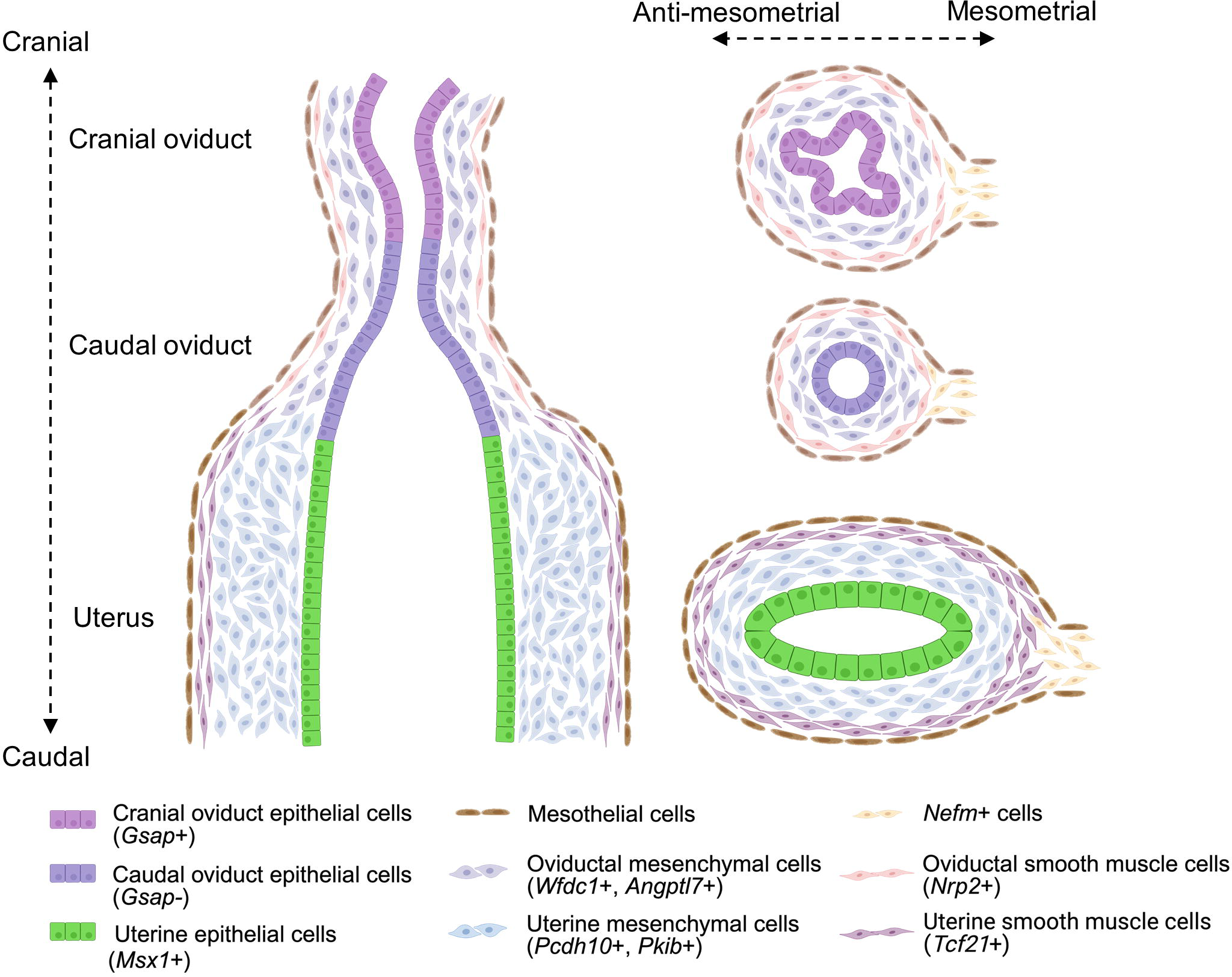
Graphical summary.

### Region-specific transcriptional signatures of the epithelium and mesenchyme in the upper Müllerian duct

Uncovering three epithelial subpopulations (*Gsap+* oviduct, *Gsap-* oviduct, *Msx+* uterine epithelium) along the craniocaudal axis at PND3 is an important finding in our study. Our results were consistent with the notion that the cranial and caudal oviducts were composed of two distinct lineages, respectively (57). MD epithelium at this developmental stage (PND3) is still uncommitted and possesses developmental plasticity till PND10. When recombined with mesenchyme, MD epithelium can be reprogrammed into epithelial cell types corresponding to the origin of recombined mesenchyme. For example, heterotypic recombinant tissues of cranial oviductal epithelium and caudal oviductal mesenchyme from PND3 neonates formed the typical cell composition pattern of the caudal oviduct (the majority is the secretory cell and few is the ciliated cell) (13). These observations suggest that while developmental fate of MD epithelium remains plastic, regional transcriptional differences have already arisen, which might play critical roles in MD regional patterning. Our identification of *Gsap* as the cranial oviductal epithelium marker at PND3 is noteworthy. The predominant number of epithelium in the cranial oviduct differentiate into multiciliate cells, which requires NOTCH signaling inhibition (58). GSAP functions as gamma-secretase activating protein that can switch the form of gamma-secretase away from Notch cleavages (59).

We identified *Msx1* as an uterine signature gene which encodes a homeobox protein functioning as a transcriptional repressor during embryogenesis (60). Previous studies have shown that MSX1 expression is detected not only in uterine epithelium but also vaginal epithelium (36). However, MSX1 expression in the vagina occurred only within the first week of postnatal development (36). Because vaginal epithelium can be elicited by uterine mesenchyme to become uterine epithelium only during the first week of postnatal development, *Msx1* was proposed to play a role in maintaining developmental responsiveness to become uterine epithelium (36). In the mature uterus, *Msx1 and Msx2* function redundantly in establishing epithelial polarity and receptivity for embryo implantation (61, 62). Therefore, these previous results along with our observations cement the notion that *Msx1* is a master regulator of various aspects of uterine epithelial development and function.

Although our dataset was powerful for comparing oviductal and uterine epithelium, it contained a relatively low number of uterine epithelia, which prevented us from further probing uterine epithelial heterogeneity at PND3. Single-cell mRNA profiling of neonatal mouse uterine epithelium at a relatively larger scale have been performed and revealed a varied number of epithelial subtypes depending on the developmental stage (63–65). These datasets together provide a wealth of information on intrinsic regulatory and signaling factors for regulating upper MD epithelial regionalization and postnatal uterine differentiation.

Our dataset also revealed distinct gene signatures for oviductal and uterine mesenchymes and their derived smooth muscles. We used region-specific Hox genes to distinguish oviductal and uterine mesenchymes at PND3 (7, 8). Since region-specific *Hox* genes are established before birth (66), these gene differences observed at PND3 might result from transcriptional regulation of region-specific *Hox* genes. Smooth muscle can be a mechanical sculptor for governing epithelial morphogenesis (67, 68). Those differential gene expression between oviductal and uterine smooth muscles might be implicated in regulating epithelial morphogenesis of these two organs: oviductal epithelia undergoes folding in the luminal side along with tubal looping while uterine epithelia are about developing epithelial budding into the surrounding stroma for forming epithelial glands. A transcription factor *Tcf21* is specifically expressed in the uterine smooth muscle. It has been reported that *Tcf21* in the mesenchyme is crucial for kidney morphogenesis (69) thus it might be interesting to determine its potential role in uterine development.

### Decoding organ-specific mesenchymal factors that potentially determine the fate of oviductal and uterine epithelia

The specific mesenchyme-derived factor(s) for directing fate decisions of oviductal or uterine epithelia *in vivo* has yet been identified. A previous study proposed that a BMP antagonist, follistatin-like-1 (*Fstl1*), was the diffusible mesenchymal factor for transforming nascent epithelial cells to oviductal ciliated cells in vitro mouse oviductal cell lines (70). However, *Fstl1* was not restricted to the oviduct in our dataset, consistent with its reported expression in the uterine region (70). In addition, *Fstl1* knockout mice died from defective lung development shortly after birth without any reports of reproductive tract abnormalities (71). The recent development of *Fstl1-flox* allele has made it feasible to determine its organ-specific role in oviductal epithelial differentiation in vivo (71, 72).

Our identifications of *Inhba* and *Gdf7* as oviduct and uterus specific mesenchymal secreted factors are intriguing. Both ligands belong to TGF-β superfamily. *Inhba* encodes inhibin βA, an essential subunit of activins and inhibin A (antagonist of activins) (73). *Inhba* is expressed in the mesonephric mesenchyme before sexual differentiation and exhibits craniocaudal expression gradients in early mesonephros (74). *Gdf7* encodes growth differentiation factor 7 and its expression is enriched in uterine stromal cells (75). In males, *Inhba* and *Gdf7* are critical for epithelial morphogenesis and differentiation of cranial and caudal Wolffian duct (primordium of the male reproductive tract), respectively (76, 77). Therefore, their significant roles in cranial and caudal MD development in females are worthy of future investigations.

### A novel mesenchymal subpopulation at the mesometrial pole

An advantage of single cell RNA-seq profiling of multicellular tissues is its capability of identifying rare cell types or subtypes that have been overlooked in the bulk tissue studies. Here, we discovered a neurofilament-positive (*Nefm*+ and *Nefl*+) mesenchymal subpopulation. As major cytoskeletal components in neuron cells, neurofilaments have been detected only in two non-neuron cells: the embryonic renal mesenchyme (78) and adrenal progenitor cells (29). Therefore, the expression of neurofilaments in one specific mesenchymal subpopulation of the upper MD is quite unexpected. In addition, *Nefm+* subpopulation resided in a unique localization, the mesometrial pole of both the oviduct and uterus. It has been noticed that the neural crest cells invade into the mesometrial pole of the Müllerian duct (79). Also, during sexual differentiation, Wolffian ducts degenerate in females however the mesenchyme surrounding Wolffian ducts is maintained and contributes to mesenchymal cells at the mesometrial side of the uterus (56). These observations suggest that *Nefm+* mesenchymal subpopulation might derive from neural crest cells and/or Wolffian duct mesenchyme.

### Extend our understanding of the less-studied mesothelial cells

Mesothelial cells have been implicated in female reproductive tract diseases, such as ectopic pregnancy and endometriosis (80, 81). In contrast to the MD epithelium lining the inside lumen, mesothelial cells form the single epithelial layer outside the female reproductive tract. Despite the opposite lining positions of mesothelial cells and MD epithelium, they have a common ancestor, coelomic epithelium (1). Coelomic epithelium is the primitive mesothelial cell lining the outside surface of the mesonephros. During early development, coelomic epithelium undergoes specification and invagination to establish the cranial portion of MD, which then extend caudally to form the entire MD epithelial tube (1, 82). We observed gene expression similarity and differences between epithelium and mesothelium, consistent with the notion that they are two lineages derived from the same progenitor. In human fallopian tubes (the oviduct), mesothelial cells were distinguished with two markers *CALB2* and *LRRN4* (83), which are different from the mesothelium marker *Msln* used in our mouse study. *MSLN* was instead enriched in human oviductal secretory cells (83–85), indicating species differences in mesothelium gene signatures. We also observed gene expression differences between uterine and oviductal mesothelial cells. During postnatal development, the oviduct becomes elongated and coiled while the uterus remains a straight tube with increased diameter. As the cellular layer lining the organ surface, mesothelial cells presumably need to undergo differential gene programs and morphological changes. For example, oviductal mesothelium had higher expression of *Igf1* which has been reported to enhance growth of oviductal organoids (86).

In summary, using scRNA-seq, we have dissected region-specific gene signatures of the epithelium and mesenchyme and deduced ligand-receptor mediated mesenchymal-epithelial interactions in the upper Müllerian duct at a critical developmental stage. We have discovered a unique mesenchymal subpopulation expressing neurofilaments and provided further understanding of pericyte and mesothelium in the oviduct and uterus. However, there are several limitations of this study. Single cell transcriptomic profiling of early or later developmental stages is absent for generating trajectory of epithelial and mesenchymal lineage progression. The role of specific subpopulations, signature genes or paracrine factors needs further confirmation by cell or gene deletions. Therefore, future studies would be necessary for comprehensively elucidating functional gene regulatory network and cell lineage differentiation in Müllerian duct development.

## Methods

### Mice

CD1 (ICR, Strain Code: 022) mice were purchased from Charles River. The day of birth was determined by pup delivery and designated as postnatal day 0 (PND0). All mouse procedures were approved by the University of Wisconsin-Madison (UW-Madison) Animal Care and Use Committees and were in compliance with UW-Madison approved animal study proposals and public laws.

### Tissue dissociation and preparation of single-cell suspension

The upper female reproductive tracts from 7 female neonates were collected freshly on PND3, when the junction between the uterus and oviduct, known as the uterotubal junction, is formed (2, 87). The connective tissues were carefully removed using a 27G needle in a petri dish with cool 1x PBS. Tracts from 3 and 4 neonatal females were pooled as two biological replicates (N=2), respectively. Oviducts and uteri were then separated at the uterotubal junction (Fig. 1A), generating four groups: oviducts (A1 and A2) and uteri (B1 and B2). Oviducts or uteri in each group were fragmented into small pieces with a 27 G needle and then transferred into a 1.5 mL tube with 250 µL dissociation media [0.04% BSA (DOT Scientific, DSA30075-25), 1.2 U/mL Dispase II (Sigma, D4693-1G), 1 mg/mL Collagenase B (Sigma, 11088807001), 5 U/mL DNase I (Sigma, DN25-100MG) in 1 mL of 1× PBS (Gibco, 14040-133)]. Tissues were dissociated at 37 ⁰C for 20 min on an orbital shaker (500 rpm). All four samples were pipetted up and down every 10 min with a 200 µL pipette until there were no observable pieces of tissue by naked eyes. The dissociation enzyme reactions were quenched by adding 1 µL 0.5 M EDTA. Media was removed by centrifugation at 500 g for 5 min at 4 ⁰C, the supernatant discarded, and pellet resuspend in 500 µL 1x PBS+0.04% BSA. Samples were transferred to 5 mL falcon tube while passing through a 35 µm cell strainer (Corning, 352235). Suspended cells were maintained on ice and sent to the Biotechnology Center at the UW-Madison for quality control (cell viability 76% (A1), 83% (A2), 77% (B1), 81% (B2)) and proceeded to 10x Genomics Single Cell mRNA Sequencing protocols.

### Single cell RNA-seq (scRNA-seq) library preparation, sequencing and alignments

Single cell sequencing libraries were prepared using Chromium Single Cell 3’ v3.1 Reagents Kits (10X Genomics) according to manufacturer’s instructions and sequenced on a NovaSeq 6000 platform using a S1 2 x 50bp flowcell. Raw data fastq files from the Illumina NovaSeq 6000 platform were aligned using default 10x Cell Ranger parameters.

### Single-cell data analysis using Seurat R package (V4.2.3)(88)

Cells with 200 < nFeature_RNA (genes per cell) < 8000, percent.mt (mitochondrial genes) < 20, and percent.Hba & Hbb (blood cell genes) < 0.5 were used for downstream analyses. Genes expressed in fewer than 3 cells were excluded from downstream analyses. The data was then normalized using a global-scaling normalization method “LogNormalize” with default setting. Highly variable features were determined by employing FindVariableFeatures function to return 2,000 features per dataset. The top 2000 highest variable features were used for the principal component analysis (PCA) and the optimal number (10 PCs) of PCA components was determined by the Elbow procedure. Single cells were clustered by the K-nearest neighbor (KNN) graph algorithm in PCA space; the dimension was reduced by the Uniform Manifold Approximation and Projection (UMAP) to visualize cell clusters. Marker genes for each cluster were identified by FindAllMarkers function (genes were detected in a minimum of 25% cells, and logFC threshold was set at 0.25). The cell type identity for each cluster was manually assigned based on established cell markers. Differentially expressed genes between clusters/subclusters were identified using the Wilcoxon rank-sum test under the Findmarkers function and then filtered by setting the adjusted p value < 0.05 and the absolute log2FC > 0.3.

The epithelial and mesenchymal clusters were isolated by the Subset function and then prefiltered, renormalized and reclustered with the following cutoffs and parameters: mesenchymal cluster: 200 < nFeature_RNA < 7000, percent.mt < 10, percent.Hba < 0.2, percent.Hbb < 0.2, UMAP resolution= 0.3; epithelial cluster: 200 < nFeature_RNA < 8000, percent.mt < 20, percent.Hba < 0.4, percent.Hbb < 0.4, UMAP resolution= 0.1. *Top2a*+ proliferating cells were excluded for eliminating the effects of different cell cycle states on reclustering and analysis of differentially expressed gene (89, 90). CellChat R package(40) was used to deduce ligand-receptor interactions between mesenchymal and epithelium cells. Average gene expression per cell cluster was calculated by using “computeCommuProb” function with the “truncatedMean” method by setting the trim value as 0.05 (5%) for Communication probabilities inferring.

### RNA-scope

The manufacturer’s protocol was followed as previously described (56). Briefly, paraffin sections were treated with antigen retrieval buffer and proteinase. Sections were hybridized with the following individual probes (*Angptl7,* Cat #: 552821; *Wfdc1,* Cat #: 471331; *Tcf21,* Cat #: 508661; *Nefm,* Cat #: 315611) and subjected to rinsing and signaling detection at designated temperatures.

### Immunofluorescence

Paraffin sections were subjected to antigen retrieval (VECTOR, H-3300) using a microwave oven. After washing with PBST (1x PBS with 0.1% Triton X-100), sections were incubated in a blocking buffer (5% normal donkey serum in PBST) for 1 hour and then incubated with a primary antibody in blocking buffer overnight at 4 °C. On the next day, sections were washed three times with PBST and incubated with secondary antibodies for 1 hour at room temperature and counterstained with DAPI (Thermo Scientific, 62248,1:1000). The following primary and secondary antibodies were used for immunofluorescence staining: Goat anti-human MSX1 Met1 Thr165 (R&D, AF5045, 1:200), Donkey anti-Goat IgG (H+L) Cross-Adsorbed Secondary Antibody, Alexa Fluor™ 488 (Invitrogen, A11055, 1:200).

## Supporting information

Code for ScRNA-seq analysis_V1

## Acknowledgments

We are thankful to Animal Care Staff at the School of Veterinary Medicine for taking care of our mice. The author(s) utilized the University of Wisconsin – Madison Biotechnology Center’s Gene Expression Center Core Facility (Research Resource Identifier - RRID:SCR_017757) for scRNA library preparation and the DNA Sequencing Facility (RRID:SCR_017759) for sequencing. We appreciate Dr. Jayshree Samanta’s suggestions on neuron cell and Dr. Shuo Xiao at Rutgers University for his help with our initial data analysis.

## Author Contributions

SJ and FZ conceived the study; FZ supervised the project; SJ performed all experiments and analyzed the data. SJ and FZ wrote the manuscript. All authors read and approved the final manuscript.

## Data and materials availability

All data are available in the main text or the supplementary materials upon reasonable request. scRNA-seq data have been deposited in the GEO database under the accession code GSE251724.

---

**Figure S1.**
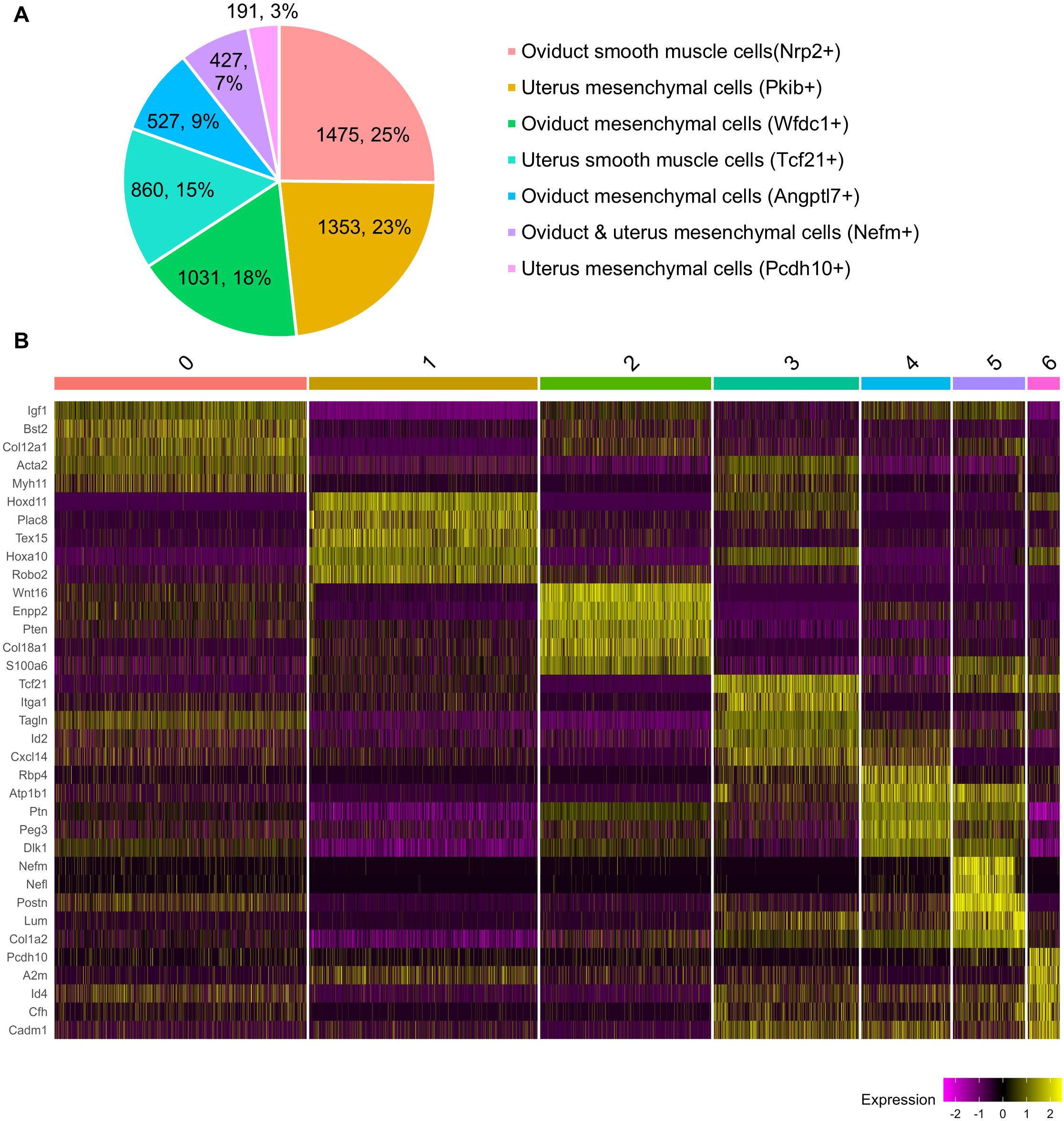
Composition of mesenchymal subpopulations in PND3 oviduct and uterus. (A) Pie chart representing mesenchymal compositions and percentages of each mesenchymal subpopulation. (B) Heatmap of top five marker genes for each cluster. The color bar for each cluster corresponds to the colors in the pie chart of Fig S1A.

**Figure S2.**
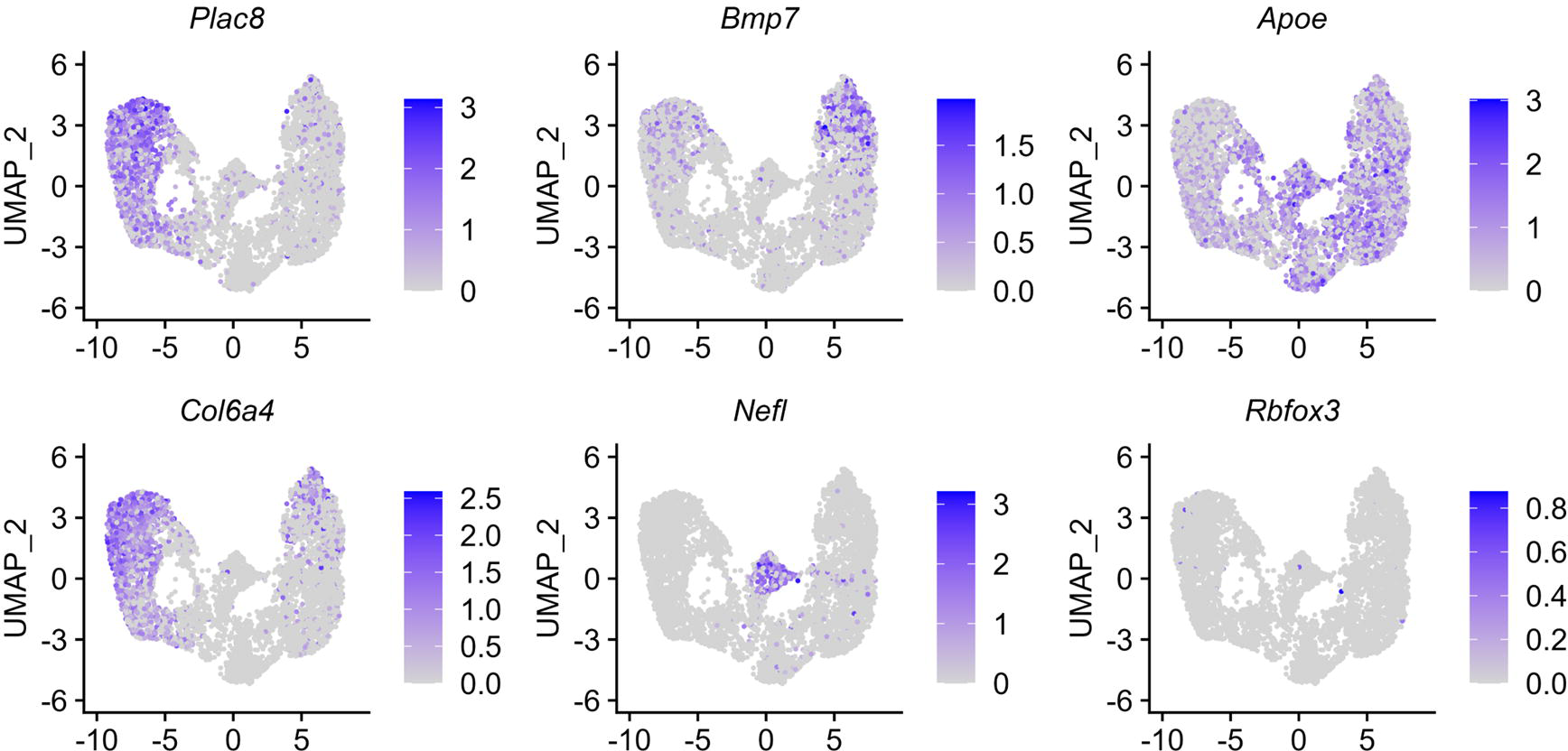
Expression of representative genes in the UMAP of the mesenchymal population.

**Figure S3.**
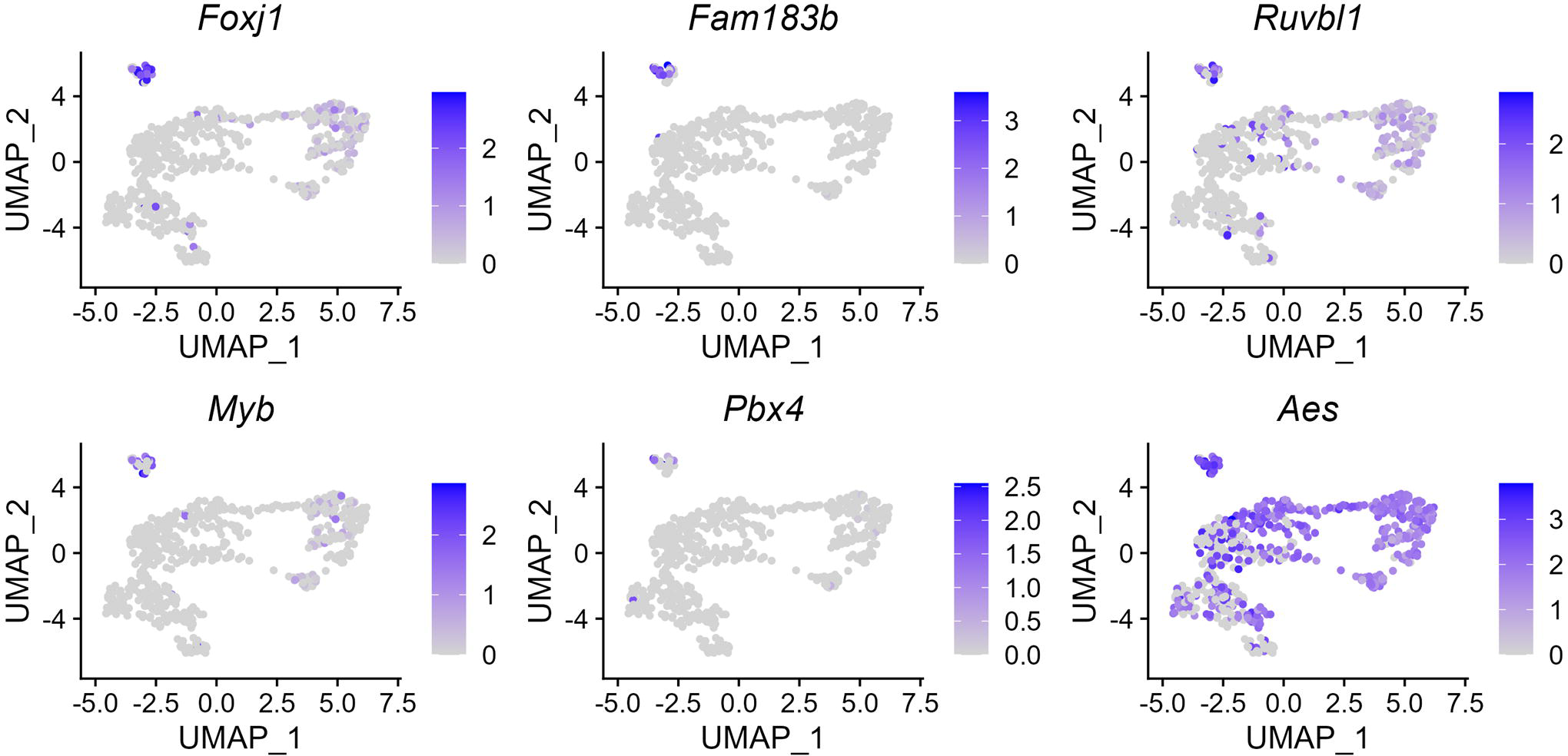
Expression of multiciliogenesis related genes in the UMAP of oviductal epithelia at PND3.

**Figure S4.**
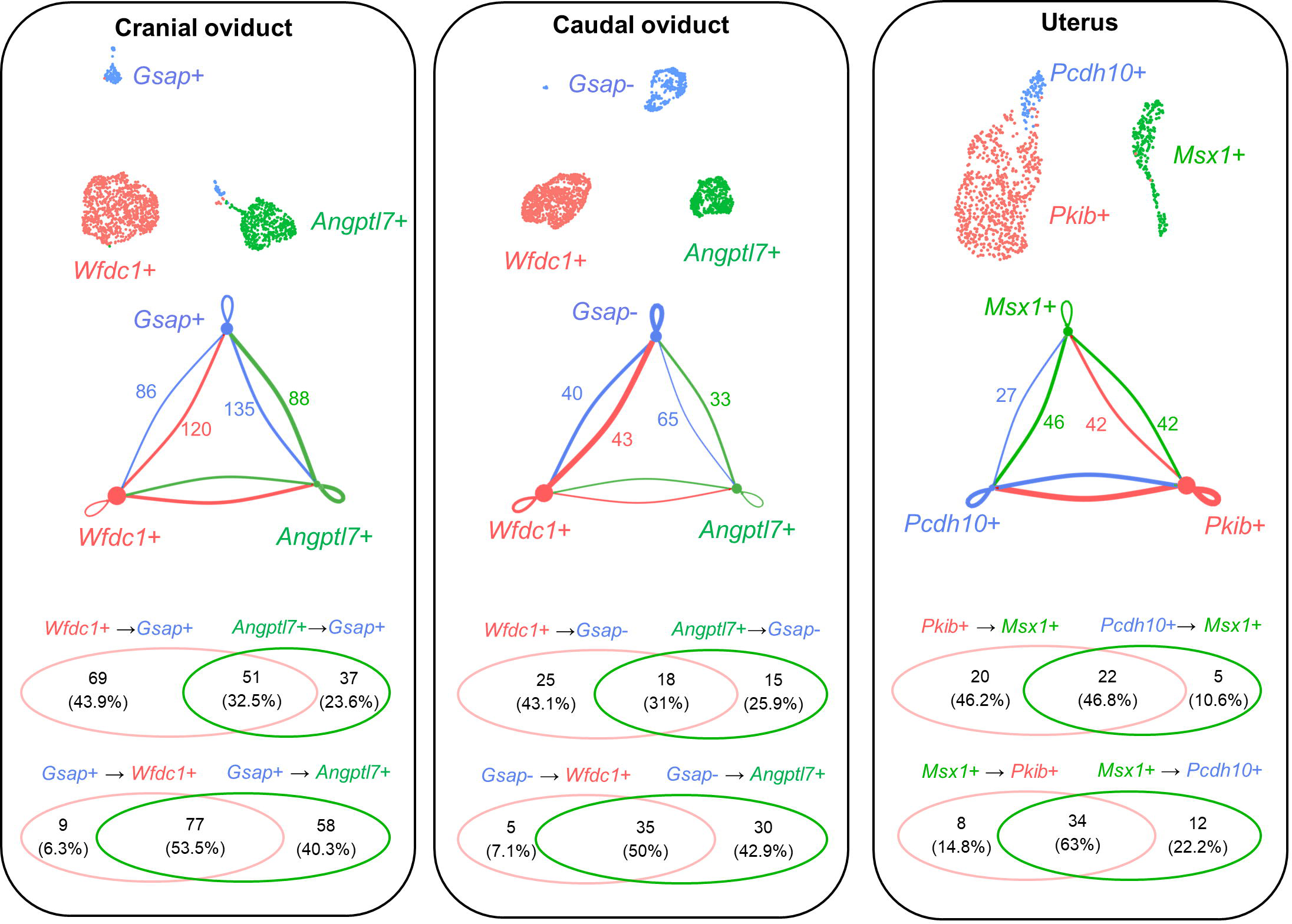
Identification of ligand-receptor pairs mediating mesenchyme-epithelium interactions in the cranial oviduct, the caudal oviduct, and the uterus at PND3. First, the number of ligand-receptor pairs mediating the crosstalk between region-specific epithelium and organ-specific mesenchymal subpopulations was shown. Then, venn diagrams showed the overlap between the lists of ligand-receptor pairs sent from two mesenchymal subpopulations to epithelium or sent from the epithelium to the two mesenchymal subpopulations. All the ligands were consolidated and selected for generating Fig 4 based on the following criteria: the knockout mice had reproductive phenotypes and/or embryonic lethality based on Mouse Genome Informatics (MGI), and ligands are enriched only in either the oviduct or uterus. Please see all ligand-pair pairs in the supplementary files 7-12.

